# Human cytomegalovirus virion delivered UL14 is a viral tropism factor for epithelial cells

**DOI:** 10.64898/2026.05.18.725932

**Authors:** Meghan F. Carter, Liz A. Kurtz, Mary Root, Eain A. Murphy

## Abstract

Infection with Human Cytomegalovirus (HCMV) can result in a significant burden of disease in those that are immunocompromised or immunonaïve. HCMV encodes a repertoire of glycoproteins that facilitate its extensive viral tropism, some of which remain to be characterized. Currently, there is no effective vaccine or cure for HCMV, therefore emphasizing the need to identify viral proteins of critical function. UL14 was selected as an open reading frame of interest due to its high scoring on an in-silico prediction algorithm, as well as its conservation amongst CMVs. Our goal was to elucidate the function of this uncharacterized viral open reading frame.

We hypothesized that UL14 functions in the establishment of infection in epithelial cells, due to its predicted structural similarity to UL141. This study demonstrates that HCMV UL14 is a glycosylated viral protein packaged with the virion. Importantly, the deletion of UL14 resulted in a significant reduction of viral growth in epithelial cells, whereas no growth defect was observed in fibroblasts. Mechanistically, we found this defect to be a result of post entry, pre-IE transcription in the establishment of infection, consistent with a defect endosomal escape. Taken together, our results suggest that UL14 functions in the establishment of infection in an epithelial cell-specific manner and may be a novel target for future vaccines or antiviral therapies.

**Author Summary:** HCMV is found in a wide variety of human cells during the course of viral infection. As such, HCMV encodes several glycoprotein complexes that dictate tropism. In this work we report the identification of a novel glycoprotein, UL14, that is involved in establishing productive infections of epithelial cells, a common site of HCMV induced disease. We report that deletion of UL14 from the viral genome impacts its ability to infect ARPE19 cells at a stage indicative of viral events post viral entry but prior to viral transcriptional activation. Further, trans complementation of UL14 by expansion of mutant virus in cells expressing the viral glycoprotein, restore viral infectivity suggesting that UL14 mediates events early in viral infection. Importantly, the characterization of this viral envelope protein provides key insights into viral tropism and identifies a novel target for vaccine design and antiviral therapies.

## INTRODUCTION

Human cytomegalovirus (HCMV) is a large beta herpes virus with high seroprevalence in the global population (Zuhair et al., 2019). As with each of the herpesvirus family members, HCMV establishes lifelong infections in a human host with recurrent activation throughout their lifetime. Individuals who are immune competent often present with no outward symptoms or may suffer from mild mononucleosis-like symptoms upon infection. Potent adaptive immune surveillance is critical for maintaining spontaneous reactivation of the virus asymptomatic. However, in individuals that are immunosuppressed such as cancer patients, transplant recipients, and those living with HIV, reactivation of the virus can cause severe complications such as end organ disease and potentially death (Plotkin & Boppana, 2019). HCMV is problematic in the immunonaïve population (i.e. neonates) as well, where it is recognized as the leading viral cause of congenital birth defects in the United States (Lantos et al., 2018). Congenital infection can lead to deafness, microencephaly, and developmental delays amongst other sequelae. There are currently no licensed vaccines against HCMV yet there are several commercially available direct acting antivirals, including ganciclovir, letermovir, cidofovir, maribavir, and foscarnet (Permar et al., 2025). However, the prolonged use of these drugs is limited due to toxicity, cross-reactivity, and the emergence of drug-resistant strains. Thus, underscoring the necessity to better understand viral tropism and modalities of entry, in hopes of one day developing a safe and effective vaccine.

The ∼236 kbp viral genome of HCMV is predicted to encode >250 open reading frames (ORFs)(Murphy et al., 2003), many of which remain to be characterized. These understudied proteins include predicted viral glycoproteins that facilitate multiple aspects of the viral lifecycle, including the broad viral tropism of HCMV. The currently well-characterized HCMV glycoproteins include gH, gL, gO, gM, gN, UL128, UL130, UL131A, UL116, UL141, and gB (Caló et al., 2016; Chandramouli et al., 2017; Ciferri et al., 2015; Kari & Gehrz, 1993; Norris et al., 2025; Siddiquey et al., 2021; Vezzani et al., 2021; Wang & Shenk, 2005). This subset of glycoproteins forms various complexes within the mature virion to facilitate cell-specific viral entry through distinct mechanisms including; membrane attachment, fusion, endocytosis, and micropinocytosis. In fibroblasts, the primary route of infection is membrane fusion, whereas epithelial and endothelial cells are predominantly infected via pH-dependent receptor mediated endocytosis (Nguyen & Kamil, 2018). The gM/gN complex binds heparin sulfate proteoglycans, acting as a tethering mechanism to the host cell, allowing for viral docking and proximity to host receptors (Kari & Gehrz, 1993). The gB homotrimer acts as the viral fusogen and is essential for viral entry; gB has also been implicated in EGFR engagement in monocytes. The remaining complexes dictate tropism and route of entry and consist of gH as the framework for the other glycoproteins. The gH/gL/gO trimer is essential for fibroblast entry where its cognate receptor is PDGFRα, although its role in entry of epithelial cells remains to be fully understood (Kabanova et al., 2016). The gH/gL/UL128/UL130/UL131A pentamer interacts with Neuropilin-2 and olfactory receptor 14I1 and is necessary for entry into endothelial cells, epithelial cells, dendritic cells, hematopoietic precursors and monocyte/macrophages (E et al., 2019; Martinez-Martin et al., 2018). Additionally, during pentamer associated entry, the Integrin/Src/Paxillin signaling pathway becomes activated, suggesting a role of integrins in this process (Chesnokova & Yurochko, 2021). Recently, a novel third gH-associated complex, GATE-3, was identified by the Kamil group (Norris et al., 2025). This gH-associated complex consists of gH/UL141/UL116, that plays a role in epithelial and endothelial cell tropism. UL141 also functions in TRAIL mediated receptor retention (Nemčovičová et al., 2013; Smith et al., 2013), suggesting that this multifunctional protein plays roles beyond tropism determination. Interestingly, UL141 is often deleted or truncated *in vitro* upon limited passage in cell culture (Cha et al., 1996; Sinzger et al., 2008).

This study focuses on the biological requirement of a previously uncharacterized viral ORF, UL14, on HCMV tropism. This ORF is predicted to encode a protein that is 343 amino acids in length that has a high probability of being expressed based on an *in-silico* bio dictionary gene finder (BDGF) algorithm which analyzes evolutionary conservation between pathogens and the human proteome (Murphy et al., 2003). HCMV UL14 maintained high amino acid conservation across different strains of HCMV including those that were serially passaged in laboratory settings. Further, the protein sequence of UL14 has homologs within evolutionarily divergent strains of herpesviruses including rhesus CMV and chimpanzee CMV(Murphy et al., 2003). While there are no reports on the function of UL14, mRNA and peptides from this ORF were identified in HCMV-wide deep sequencing and proteomic analysis (Rozman et al., 2022; Song et al., 2023a; Weekes et al., 2014).

Herein, we report that our predictive structures and *in silico* analysis suggest that this ORF encodes a type-I transmembrane protein with a glycosylated ectodomain, that shares high structural homology to HCMV encoded UL141, the main component of the GATE-3 (Norris et al., 2025). These attributes lead us to the hypothesis that UL14 functions in the establishment of infection in epithelial cells. We demonstrate that UL14 is critical for the establishment of infection in epithelial cells *in vitro.* Our data reveal UL14 mRNA and protein are expressed in fibroblasts and epithelial cells with Early/Late expression kinetics, consistent with previous studies. Furthermore, our data shows UL14 to be a glycosylated viral protein that co-localizes with the viral assembly compartment (VAC) and is packaged with the virion. Through the generation of sequential viral mutants, we demonstrate that the complete deletion of UL14 results in an epithelial cell specific growth defect, wherein we observed that the deletion of UL14 significantly reduces the expression of the critical lytic protein, IE1, when compared to wild-type infection, despite the same number of viral genomes being delivered to each cell. Additionally, when the ΔUL14 virus is trans complemented by propagation in UL14-expressing fibroblasts, wherein UL14 is incorporated into the virion but not expressed *de novo*, we observe that the envelope delivered UL14 rescues the epithelial IE1 phenotype. Mechanistically, we observe that in the absence of UL14, viral infection is halted at the endosome in epithelial cells, preventing viral uncoating and nuclear delivery. Taken together, our results suggest that UL14 functions in the early establishment of infection in an epithelial cell-specific manner.

## RESULTS

### HCMV UL14 shares predictive structural homology to HCMV UL141

UL14 is of interest due to its high amino acid conservation across clinical isolates as well as laboratory adapted strains and in chimpanzee CMV (ChCMV) and rhesus CMV (RhCMV), (**Supplemental Figure 1a & 1b**) and its mRNA and peptides from this ORF were identified in HCMV-wide deep sequencing and proteomic analysis (Rozman et al., 2022; Song et al., 2023b; Weekes et al., 2014). Due to the lack of published information regarding the structure or function of UL14, we first generated a predictive protein structure for the full-length amino acid sequence using AlphaFold 3 (Jumper et al., 2021) coupled with analysis with Signal-IP 6.0 (Teufel et al., 2022), DeepTMHMM (Hallgren et al., 2022), and Topcons (Tsirigos et al., 2015) which identified the putative domains of the protein; including an N-terminal signal peptide at amino acids 1-23 with a cleavage site between amino acids 23 and 24, an Ig-like ectodomain at amino acids 23-265, a transmembrane domain at amino acids 266-286, and a cytoplasmic tail at amino acids 287-329 **(Figure 1a**). This structure was then uploaded into the Fold Seek platform (van Kempen et al., 2024), which searches the Protein Data Bank (PDB) for homologs based on structure in addition to sequence homology. In doing so, we identified that HCMV UL141 as having structural homology to the UL14 ectodomain, with a probability of 1.0, a high sequence identity of 28%, an E-value of 5.79e-17, a TM-score indicative of global structure similarity of 0.823, and an RMSD reflective of local atomic alignment of 3.16 (**Figure 1b and 1c**). These *in silico* analysis techniques demonstrated predictive structural homology of UL14 to UL141 regarding the putative signal peptide and Ig-like domains, suggesting a potential for a shared function.

**FIGURE 1:**
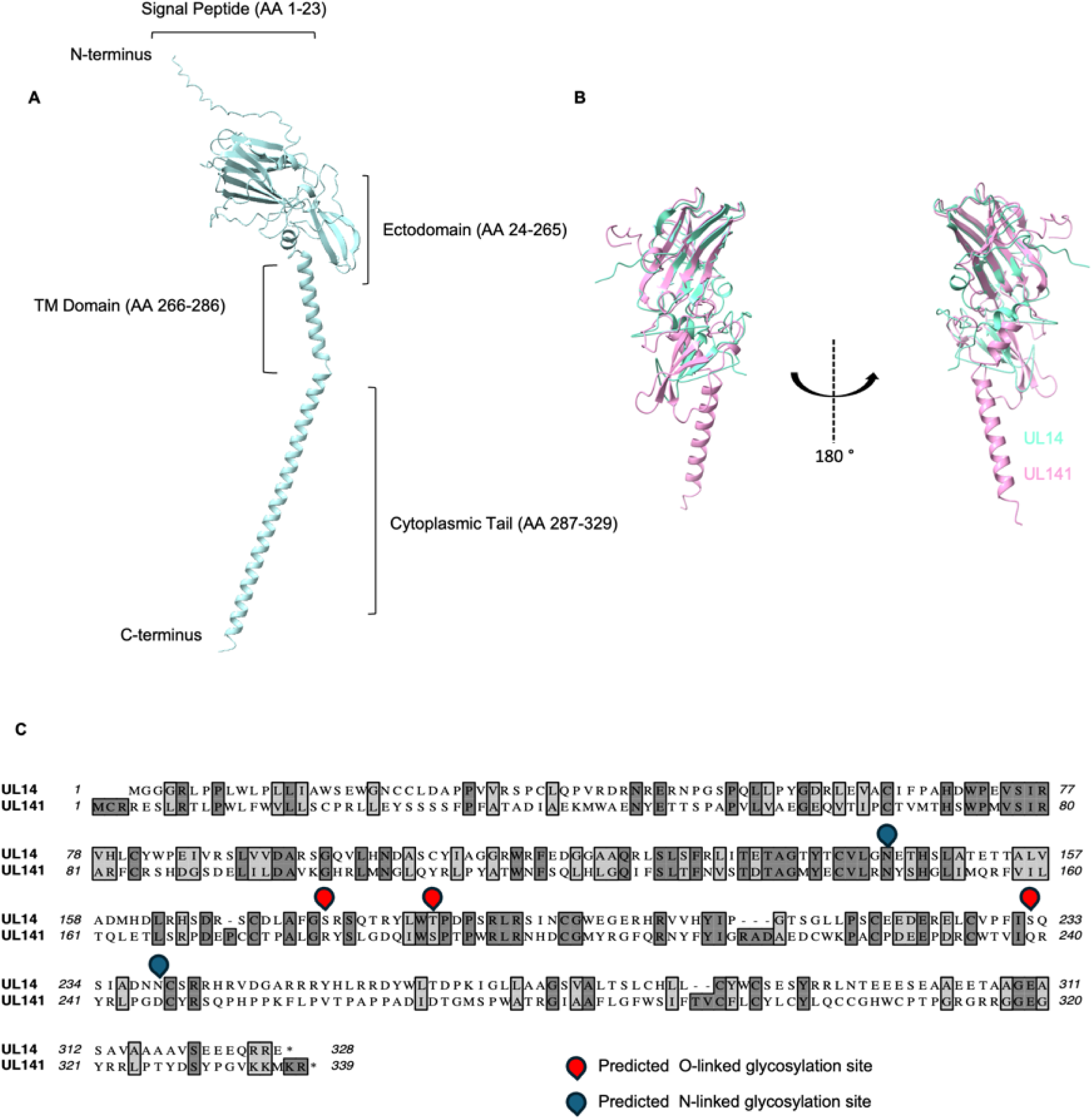
HCMV UL14 shares predictive structural homology to HCMV UL141. Alphafold predicted structure of UL14 with putative domains. **a**) Annotated UL14 sequence of putative domains according to SignalIP 6.0, DeepTMHMM, and TOPCONS. **b**) Structural homology analysis of UL14 Alphafold prediction to published proteins queried using the Foldseek platform reveals UL141 as a structural homolog. Structural rendering of UL14 ectodomain (green) overlayed with HCMV UL141 ectodomain (pink) with a Prob of 1.0, sequence ID of 28%, an E-value of 5.79e-17, a TM-score of 0.823, and an RMSD of 3.16. **c**) Clustal amino acid alignment demonstrating conservation of UL14 and UL141 with annotated UL14 predictive sites of glycosylation.

### The UL14 transcript and protein are detected in HCMV infected fibroblasts and epithelial cells

Although previous studies identified UL14 transcripts in RNA-sequencing data sets, and peptides that matched the predicted protein encoded by UL14, we wanted to confirm that in our cells of interest, the UL14 mRNA transcript and protein is expressed during lytic infection. As there is no commercially available antibody raised against UL14, we generated a Bacterial Artificial Construct (BAC) recombinant HCMV in which the carboxy terminal domain of UL14 was fused in frame with three tandem FLAG tag epitopes termed TB40/E-mCherry-UL99eGFP-UL14-3XFLAG (UL14-FLAG) (**Figure 2)**.

**FIGURE 2:**
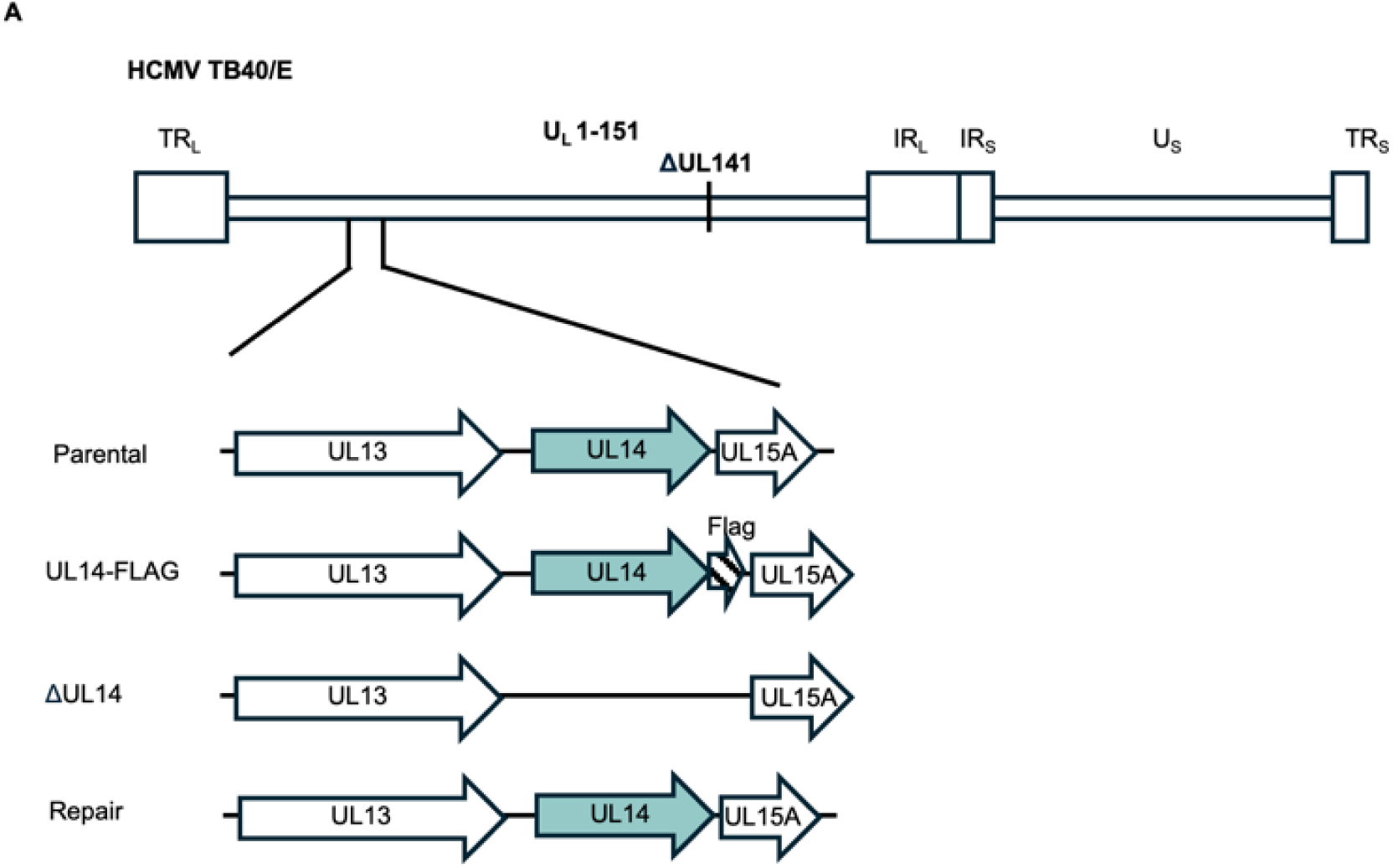
Schematic of UL14 viral mutants generated in the TB40/E background. The parental TB40/E (TB40/E-mCherry-UL99eGFP) was used to sequentially generate UL14-FLAG (TB40/E-mCherry-UL99eGFP-UL14-FLAG), ΔUL14 (TB40/E-mCherry-UL99eGFP-UL14), and the repair virus using BAC recombineering.

To first monitor UL14 transcription, we infected highly lytic fibroblasts (Nuff-1 cells) with HCMV (strain TB40/E-mCherry-UL99eGFP) (WT) at a multiplicity of infection (MOI) of 1. RNA was collected at 4 and 24 hours post infection and assessed for expression of UL14 and GAPDH transcripts by RT-qPCR. The UL14 transcript was detected at both 4 hours, and 24 hours post infection (**Figure 3a**) indicating that the promoter of UL14 was transcriptionally active during lytic infections. To evaluate the accumulation of UL14 protein expression by immunoblot, we used UL14-FLAG to infect primary fibroblast cells (Nuff-1 cells) or epithelial cells (ARPE-19 cells) at an MOI of 1, and cell lysates were collected across a time course of 96 hours. Lysates were then probed using anti-M2 to detect the FLAG peptide or tubulin as a loading control. UL14-FLAG was detectable by western blot as early as 12 hours in fibroblasts (**Figure 3b**) and 24 hours in epithelial cells (**Figure 3c**) suggesting that the protein is expressed and accumulates to significant levels at late times during infection.

**FIGURE 3:**
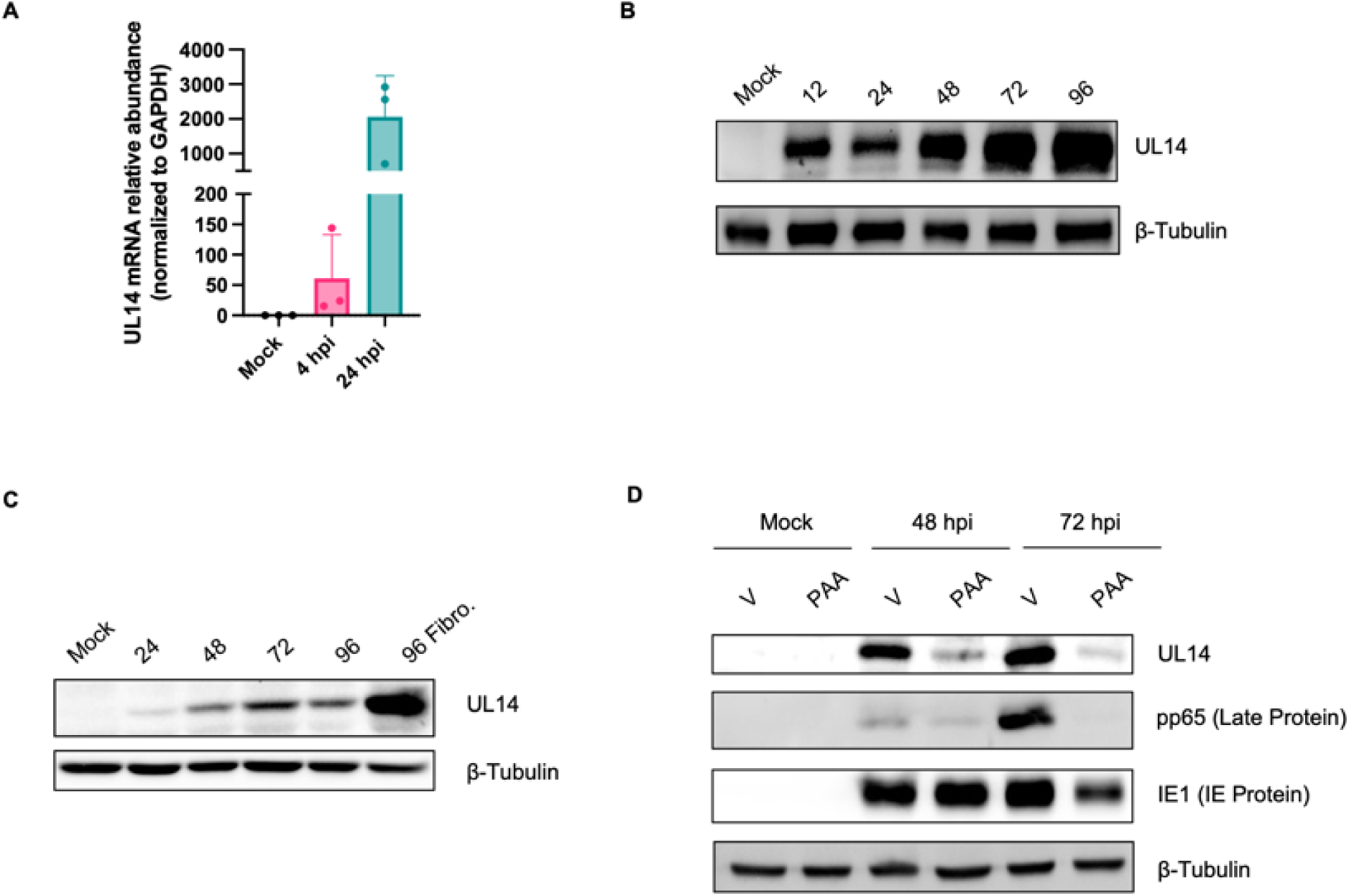
The UL14 transcript and protein can be readily detected in fibroblasts and epithelial cells. **a**) Nuff-1 cells were infected at an MOI of 1.0 TCID_50_/cell using the TB40/E-mCherry-UL99eGFP virus. RNA was collected at 4 and 24 hours and UL14 transcripts were quantified by qPCR and normalized to GAPDH (*N = 3)* **b)** Primary fibroblasts (Nuff-1) or **c**) Epithelial cells (ARPE-19) were infected at a multiplicity of 3.0 TCID_50_/cell using the TB40/E-mCherry-UL99eGFP-UL14-FLAG virus. Cell lysates were collected across a time course of 96 hours and probed by western blot for the UL14 protein using anti-flag; β-Tubulin was used as the loading control. Representative blots are shown (*N = 3*). **d**) Primary fibroblasts were infected at an MOI of 1.0 TCID_50_/cell and treated with vehicle or PAA (300ug/mL) and harvested at 48 and 72 hours. Cell lysates were probed by western blot for UL14, pp65, IE1, and β-Tubulin. (*N = 3*)

Additionally, we wanted to confirm the expression kinetics of this protein, as UL14 peptides were previously identified in a proteomic screen as being of the *Early* temporal class (Rozman et al., 2022; Weekes et al., 2014). We pre-treated cells with Phosphonoacetic Acid (PAA), a viral DNA polymerase inhibitor which blocks viral DNA replication and therefore allows us to delineate between early gene expression and late gene expression. Fibroblast cells were infected at an MOI of 1 with the UL14-FLAG virus, treated with vehicle (DMSO) or PAA (300ug/mL), and then cell lysates were collected at 48 and 72 hours post infection. Lysates were then probed by immunoblot analysis for expression of UL14, the true Late protein, pp65 (as control for PAA treatment) and β-tubulin. IE1 protein expression served as a control as it does not require *de novo* viral DNA synthesis and should not be significantly impacted by PAA, and β-tubulin protein assessment served as a loading control. We observed that the expression of UL14 protein was present but reduced upon treatment with PAA, suggesting that UL14 is expressed with Early/Late kinetics (**Figure 3d**), consistent with previous publications in which peptides and mRNA from UL14 were identified in proteomic and deep-sequencing screens as having Early kinetics of expression. These results show that the UL14 transcript and protein are expressed during lytic infections of both highly permissive fibroblasts and clinically relevant epithelial cells and accumulate at times consistent with the initiation of VAC formation.

### UL14 is a viral glycoprotein that colocalizes with the viral assembly complex and is packaged in the virion

As UL14 *in silico* analysis revealed high predictive amino acid structural conservation with UL141, an envelope encoded glycoprotein involved in epithelial tropism, we wanted to evaluate the glycosylation status of UL14. We utilized two algorithms, NetNGlyc-1.0 and NetOGlyc-4.0, to determine the predictive glycosylation status of UL14. These tools identified amino acid position N144 (-<u>N</u>ETH-) and N239 (-<u>N</u>CRS-) to be possible N-linked glycosylation sites, with a potential of 0.6 and 0.4, respectively. Additionally, amino acid residues S176, T185, and S232 were predicted to be O-linked glycosylation sites (**Figure 1c)**. To experimentally validate the N-linked glycosylation status, we infected fibroblasts with the UL14-FLAG virus (MOI of 1) and harvested whole cell lysates at 120 post-infection. Lysates were then treated with PNGase F which removes all N-linked carbohydrates, EndoH which removes high mannose and some hybrid N-linked carbohydrates, or lysates were untreated. We then monitored shifts in the UL14 migration status using the M2 antibody by immunoblot analysis with β-tubulin expression serving as a loading control. A faster migrating FLAG specific protein of an approximate 14 kDa shift from 50 kDa to 36 kDa was observed in both treated lysates when compared to untreated lysates, of which corresponds to the predicted unmodified size of UL14 which is approximately 36 kDa, (**Figure 4a**) suggesting that UL14 is an N-linked glycosylated viral protein.

**FIGURE 4:**
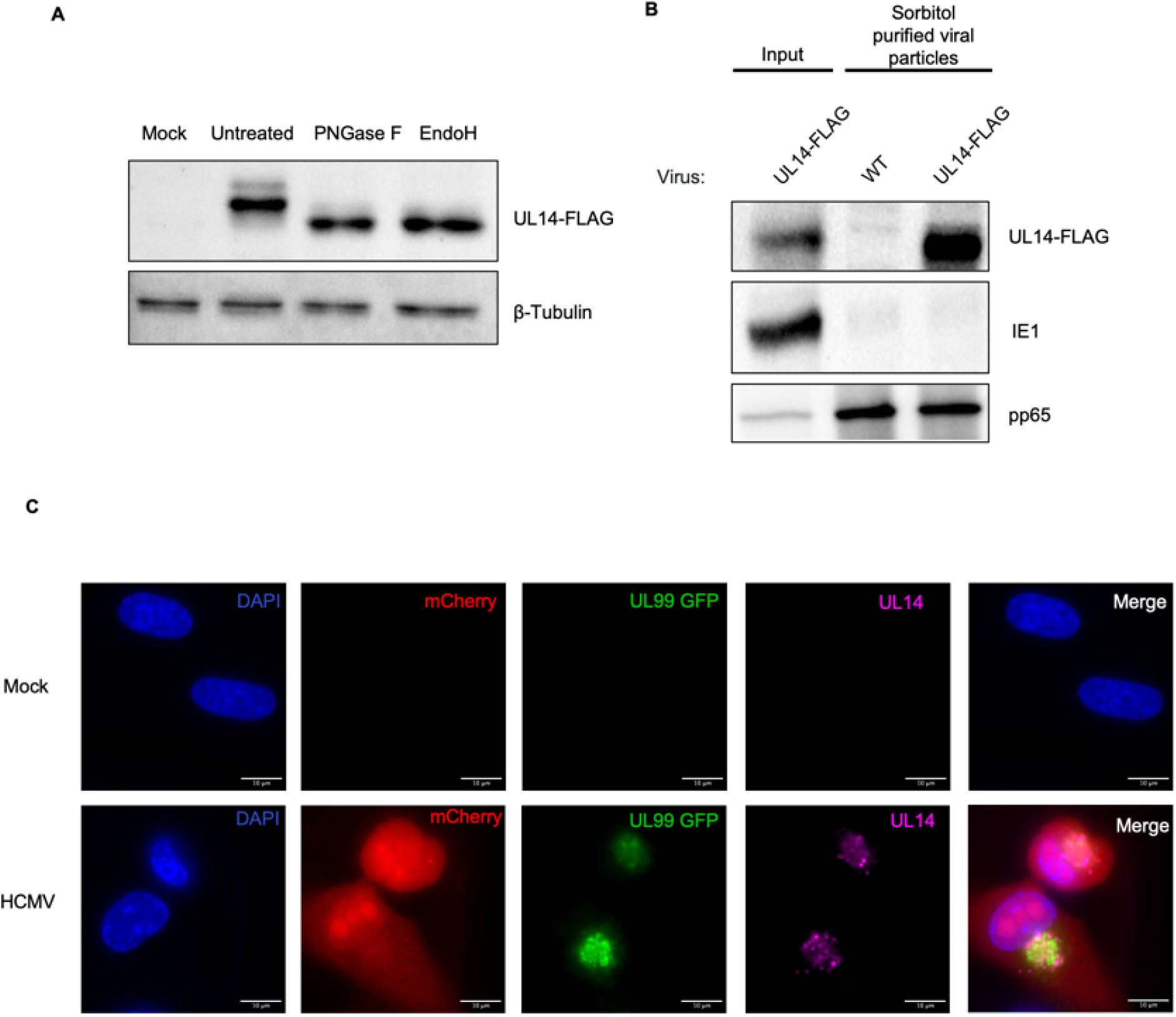
UL14 is a viral glycoprotein that colocalizes with components of the the viral assembly complex and is packaged in the virion. **a)** Fibroblast were infected at a multiplicity of 1.0 TCID_50_/cell using the TB40/E-mCherry-UL99eGFP-UL14-FLAG virus and cell lysates were harvested and treated with PNGase or EndoH at 120 hours post-infection (*N = 3)*. **b**) UL14-FLAG virus or WT virus were expanded in fibroblasts. Cell associated and cell free virus was harvested after 100% CPE was observed, by ultracentrifugation through a 20% sorbitol cushion. Viral pellets were then used for western blotting, where UL14 was probed for using anti-flag; anti-pp65 and anti-IE1 were used as controls (*N = 3*). **c**) NuFF-1 cells were infected with HCMV at an MOI of 1.0 TCID_50_/cell. Cells were fixed, stained with their respective antibodies, and imaged at 72 hours post-infection (*N=3)*.

Given that UL14 is glycosylated and has predicted structural homology to UL141, we next wanted to know if UL14 is packaged within the virus. To evaluate this, we infected low passage WT and UL14-FLAG virus in fibroblasts until they reached 100% CPE. Cell-free and cell-associated virus was then isolated using a 20% sorbitol cushion, and the viral pellet was used to generate lysates suitable for immunoblotting. We also included whole cell lysates from infected fibroblasts (input) as a positive control. The blot was then probed for UL14 using the M2 antibody and IE1 to confirm purity of the viral pellet as IE1 is not found in the virion. Further, we blotted for pp65 (UL82) as this protein is found in the tegument thus serving as a positive control for virion isolation. Our cell free lysates were negative for IE1 staining and positive for pp65 staining assuring that our purification process successfully isolated tegument proteins yet lacked proteins indicative of cell lysate contamination. Importantly, UL14 was robustly detected within the sorbitol purified viral particles **(Figure 4b)**, demonstrating that UL14 is indeed packaged within cell free virions and delivered to cells upon infection.

Given that UL14 is glycosylated and packaged with the virus, we wanted to confirm its intracellular localization using IFA. Fibroblasts were infected with the UL14-FLAG virus for 72 hours, fixed, stained, and imaged. To monitor viral infection, we monitored expression of the fluorescent protein, mCherry, which was engineered into our recombinant virus downstream of the SV40 early promoter and expressed with IE kinetics. Additionally, this virus expresses eGFP which is fused in frame to the UL99 protein which is expressed with late kinetics and localizes to the VAC. UL14 was monitored using the M2 antibody raised against the flag peptide and DAPI was used to stain the nucleus and uninfected cells were used as a staining control. Within mCherry positive infected cells, UL14 was detected in subcellular regions consistent with UL99-eGFP expression at the perinuclear site of viral assembly (**Figure 4c**). Thus, further supporting that UL14 is a glycosylated viral protein that localizes to the VAC where HCMV glycoproteins are packaged with the virion.

### UL14 is critical for growth in epithelial cells but not fibroblasts

To define the requirement of UL14 during viral infection, we generated two additional mutants to complement our UL14-FLAG virus, consisting of a complete deletion of UL14 (ΔUL14), and then a sequential repair of the complete deletion (Repair) **(Figure 2).** The generation of the Repair virus assures that any observed phenotype in deletion virus is not due to a non-specific alteration in the viral genome beyond the UL14 ORF. We then assessed the growth kinetics of these viruses over time in fibroblasts and epithelial cells. We included epithelial cells in this analysis due to the phenotype observed with a UL141 deletion where there was no growth defect in fibroblasts; yet there was a 5-fold reduction in viral progeny production of epithelial cells. Using an MOI of 1, we infected both cell types and measured viral production by TCID_50_. In fibroblasts, there was no appreciable difference in the growth kinetics of the UL14 deleted virus when compared to either the WT or Repair virus, suggesting the viral protein is expendable for lytic replication in highly permissive fibroblasts **(Figure 5a).** However, in epithelial cells, we observed a 2-log reduction in viral replication of the ΔUL14 virus, in comparison to the parental virus. Furthermore, this growth defect could be rescued by repairing the complete deletion of UL14 **(Figure 5b and Supplemental Figure 2).** These results demonstrate that UL14 plays a cell-specific role in establishing productive viral infections of epithelial cells that require endocytosis of the virus for infection compared to fibroblasts where infection occurs at plasma membrane fusion.

**FIGURE 5:**
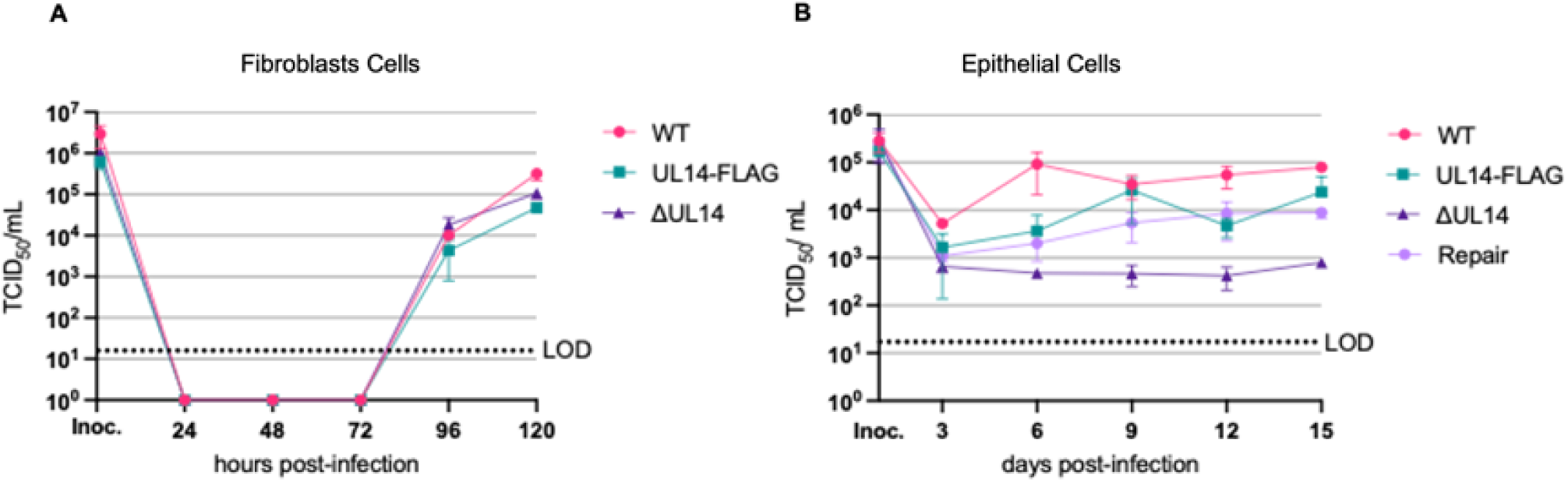
Growth kinetics of UL14 mutants in comparison to WT. **a)** Primary fibroblasts (NuFF-1) were infected with the viruses indicated at an MOI of 1.0 TCID_50_/cell. Supernatants were collected over a time course of 120 h, and then viral titers were quantified for each time point by TCID_50_ assays. **b**) Epithelial cells (ARPE-19) were infected with the indicated viruses at a multiplicity of infection of 1.0 TCID_50_/cell. Cell associated virus was collected over a time course of 15 days, and then viral titers were quantified for each time point byTCID_50_ assays. Mean ± SD is shown (*N = 3*).

### IE1 protein expression is diminished in epithelial cells in the absence of UL14

We next wanted to establish at what point during viral infection is UL14 most critical. To evaluate this, we first looked at IE1 protein expression following the infection of epithelial cells. We infected ARPE-19 cells at an MOI of 3 with WT, UL14-FLAG, ΔUL14, and the Repair virus, respectively. Lysates were then collected at 48 hours post-infection and used for immunoblotting. We probed for IE1 expression and β-tubulin staining served as a loading control. We observed that IE1 protein expression was significantly reduced when epithelial cells were infected with the ΔUL14 virus, in comparison to WT and UL14-FLAG. Importantly, this ablation of IE1 protein expression was rescued using the Repair virus **(Figure 6a and b**) assuring the defect was due to the loss of UL14 expression and not any other defect in the viral genome. These results suggest that the presence of UL14 is essential for a key step of HCMV viral infection that exists between viral attachment and that of efficient initiation of IE gene expression.

**FIGURE 6:**
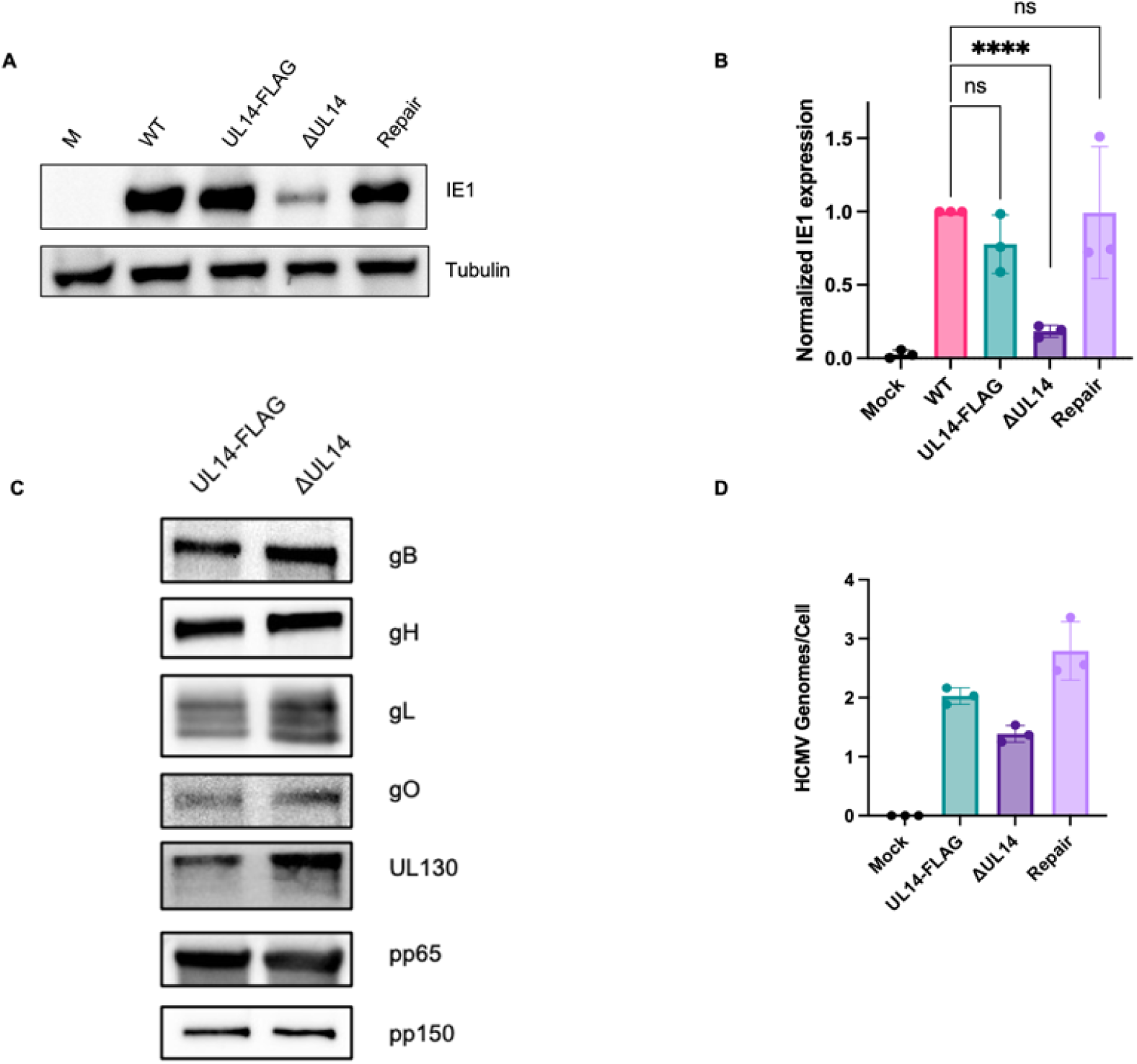
IE1 expression is diminished in epithelial cells when UL14 is deleted. Epithelial cells (ARPE-19) were infected with the respective viruses at a multiplicity of infection of 3.0 TCID_50_/cell. **a**) Cell lysates were collected at 48 hours post infection and IE1 was probed by western blot using the anti-iE1 antibody; β-Tubulin was used as the loading control. **b**) Densitometry mean ± SD is shown, statistical significance calculated using *t*-test; *** p ≤ 0.0001 (*N = 3*). **c**) Viral glycoprotein and pp65 loading of cell-free virions was determined by western blot and pp150 served as a loading control (*N = 3*). **d**) Viral and cellular DNA were quantified by qPCR using primers targeting the UL123 region of the viral genome and MDM2 targeting the human genome, respectively. Genome copy numbers were determined through the employment of a BAC that contains both UL123 and MDM2, mean ± SD is shown (*N=3*).

### Cell free virions have equivalent glycoprotein loading in the absence of UL14, when compared to parental virus

Given that the ratio of viral glycoproteins loaded onto the virion is a recognized determinant of viral tropism (Scrivano et al., 2011), we wanted to investigate if the absence of UL14 altered the glycoprotein composition of cell free virus. To evaluate this, we expanded low passage UL14-FLAG virus and ΔUL14 virus on fibroblasts until they reach 100% CPE. The viral particles were then isolated through a 20% sorbitol cushion using ultracentrifugation, and lysates were prepared for western blotting. The blots were probed for the tegument protein pp150 as a loading control, gH, gB, gO, gL, UL130, and pp65 as tegument control. Across both conditions viral glycoprotein loading and pp65 loading were equivalent (**Figure 6c**), suggesting that the epithelial cell specific infection deficit is not due to alterations in the loading of these specific glycoproteins, or tegument proteins.

### An equal number of viral genomes are being delivered to each infected cell

To determine if the reduced IE expression in epithelial cells infected with ΔUL14 was a result of a reduction in viral infectivity of ARPE19 cells, we monitored the number of HCMV genomes delivered upon infection. To this end, we infected epithelial cells at an MOI of 1 and removed unbound virus by washing cells 1 hour post infection. We then treated the cells with trypsin at 6 hours post infection to effectively remove any bound virus that was attached to the outside of the cells but failed to enter the cells. Total cellular and viral DNA were then harvested and quantified by qPCR using HCMV UL123 and human MDM2-specific primers. A standard curve with a defined number of copies for UL123 and the MDM2 gene was used to allow quantification of viral genomes per cell. We observed a near equivalent number of viral genomes of UL14-FLAG, ΔUL14 and Repair virus (1.5 to 3 viral genomes/cell) being delivered to cells upon treatment (**Figure 6d**). This result suggests that the IE1 protein accumulation deficiency observed upon infection of ARPE19 cells with ΔUL14 virus was not due to a defect in viral cell attachment and genome entry. However, the lack of IE transcription supports the possibility that the viral genomes present within nucleocapsids fail to translocate to the nucleus.

### Virion delivered UL14 rescues IE1 phenotype in epithelial cells

Given that UL14 is found to be packaged in the virion, we wanted to know if virion associated UL14 is responsible for early steps of viral infection resulting in efficient IE1 protein accumulation within ARPE19 cells. To evaluate this, we expanded low passage (P0) ΔUL14 virus within transduced fibroblasts that stably express the UL14 protein (**Figure 7a**). By replicating the ΔUL14 virus within these UL14 expressing cells, the viral glycoprotein would be available to be incorporated into newly synthesized viral particles. We confirmed that this virus, denoted UL14-Comp virus, contains trans-complemented UL14 packaged within cell free virus and could deliver virion associated UL14 to naïve cells by infecting fibroblast at a high MOI (10), harvesting whole cell lysates at 6hpi and immunoblotting for UL14 using the M2 antibody and β-tubulin as a loading control. We observed that UL14 protein could be detected in naïve cells infected with the UL14-Comp virus showing that the transduced cells allowed for packaging of UL14 in the mutant virus **(Figure 7b).**

**FIGURE 7:**
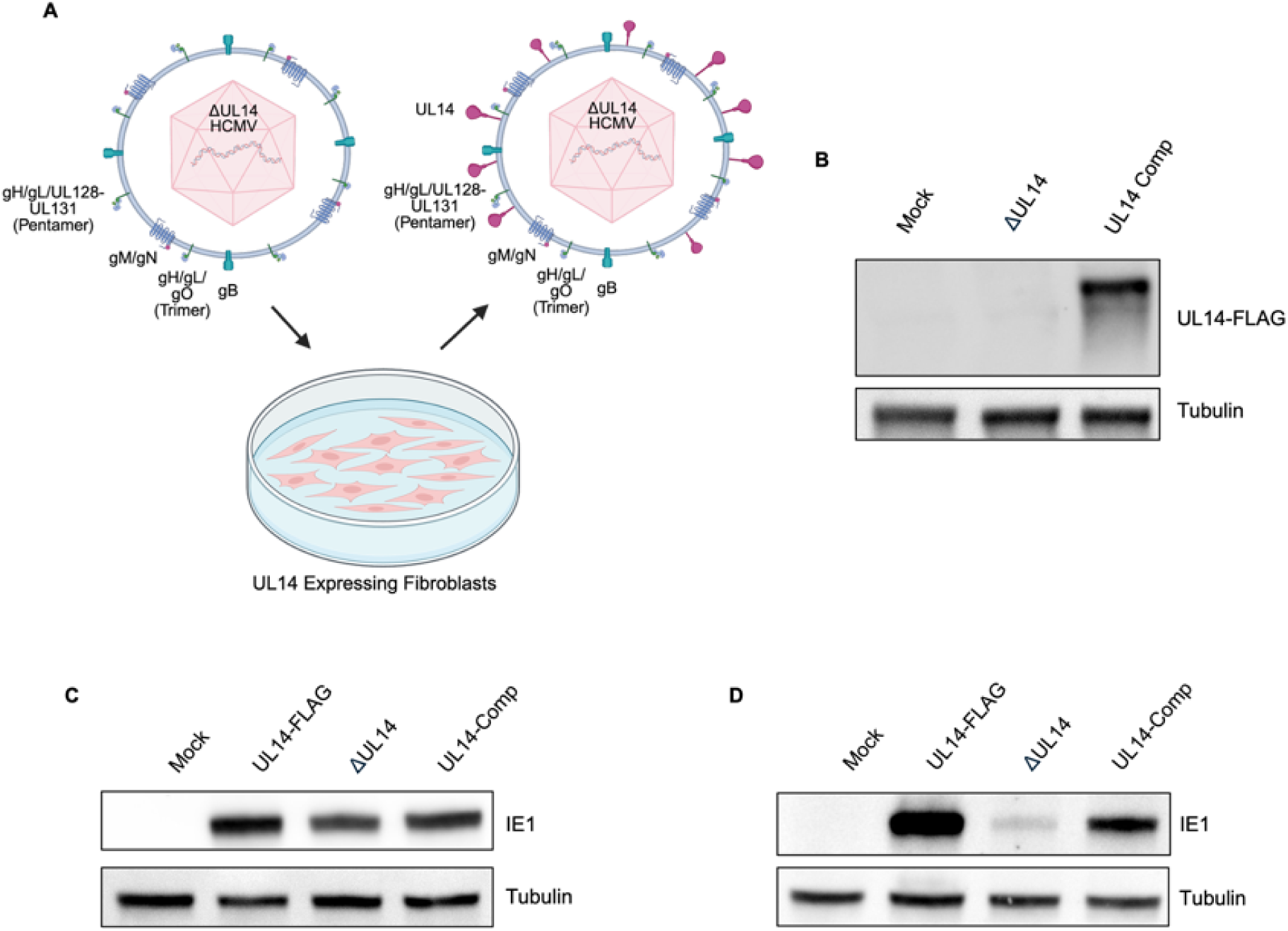
Virion delivered UL14 rescues IE1 phenotype in ARPE-19 cells. **a**) schematic of ΔUL14 virus passaged through UL14-FLAG stable expressing fibroblasts to generate UL14-Comp virus. **b**) Fibroblasts were infected with the respective viruses at an MOI of 10 TCID_50_/cell and cell lysates were harvested at 6 hours post-infection and then used for western blotting, where UL14 was probed using anti-flag and β-Tubulin was used as a control (*N = 3*). **c**) fibroblasts or **d**) epithelial cells were infected with the respective viruses at an MOI of 3.0 TCID_50_/cell. Cell lysates were collected at 48 hours post-infection and then used for western blotting. IE1 was probed for using anti-iE1 antibody and β-Tubulin was used as a loading control (*N = 3*).

Next, to determine if the UL14-Comp virus can rescue the cell type specific reduced E1 protein expression phenotype in epithelial cells, we infected fibroblasts and ARPE19 cells using the UL14-FLAG, ΔUL14 virus, and the UL14-Comp virus. Cell lysates were collected at 48 hours post infection and used for immunoblotting using antibodies specific for IE1 and β-tubulin as a loading control. As predicted, the expression of IE1 protein was not appreciably different in fibroblast infected cells as UL14 is dispensable for efficient viral infection of this cell type (**Figure 5a and 7c**). Importantly, the ΔUL14 virus infection of ARPE19 cells again showed reduced expression of IE1 protein upon infection whereas the UL14-Comp virus demonstrated increased IE1 expression in relation to the infected cells that were infected with virus that was not complemented. It is important to note that the UL14-FLAG tagged virus infection of ARPE19 cells demonstrated levels of IE1 protein that were higher than the UL14-Comp infected cells, which may be due to efficiency of UL14 loading into the UL14-Comp viral particles or due to newly synthesized virus within the UL14-FLAG infected cells that can then infect naïve ARPE19 cells (**Figure 7d).** Overall, these finding support the premise that complementation of UL14 into viral particles that lack the genomic capacity to synthesize UL14 can rescue the IE1 deficiency phenotype in epithelial cells when compared to the ΔUL14 virus and that this viral protein is necessary to mediate viral lifecycle events at prior to IE transcription in epithelial cells.

### Nuclear delivery of pp65 and endosome escape is compromised in the absence of UL14 in epithelial cells

Our observed UL14-dependent viral tropism defect was observed in ARPE19 cells but not fibroblasts. As mentioned above, viral entry is distinct between fibroblasts (fusion from without) vs. epithelial cells (pH dependent endosome fusion). Thus, it is possible that the ΔUL14 virus can enter epithelial cells but is not able to escape endosomes. To determine if viral endosome escape is being impacted by UL14, we infected ARPE19 cells with UL14-FLAG and ΔUL14 virus and monitored the nuclear accumulation of tegument protein pp65 as a surrogate marker of efficient viral release from the endosome. In this instance, pp65 is a suitable control as it is loaded equally into both viruses of interest (**Figure 6c**). Infected epithelial cells (MOI of 5) were fixed and stained at 15 hours post infection with DAPI to highlight the nucleus and anti-pp65. To visualize early endosomes, we stained for Early Endosome Antigen 1 (EEA1). In UL14-FLAG viral infections, we observed efficient pp65 delivery to the nucleus suggesting tegument release from the endosome. However, in the absence of UL14, we found that the tegument pp65 packaged in the ΔUL14 virus was not found to accumulate in the nucleus but instead was found in regions of the infected cells that stained positive with EEA1 **(Figure 8).** This result supports a model in which the IE defect observed in the ΔUL14 infected epithelial cells may be due to an inability for the virus to escape from the endosome, preventing tegument release from viral particles. This would result in a lack of genome delivery to the nucleus, thus resulting in diminished IE1 transcription.

**FIGURE 8:**
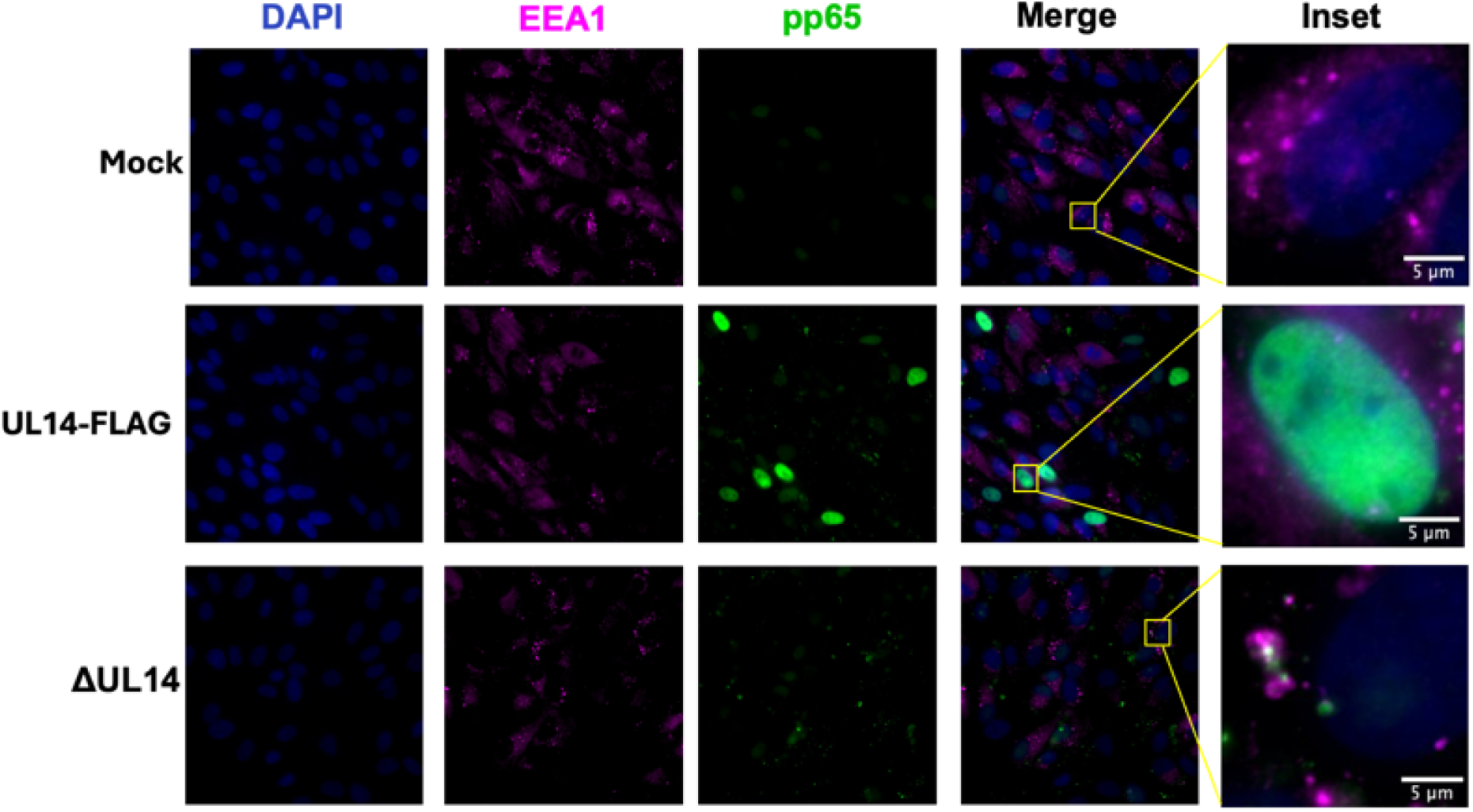
Nuclear delivery of pp65 and endosome escape is compromised in the absence of UL14 in epithelial cells. Epithelial cells were infected with the indicated viruses at an MOI of 5.0 TCID_50_/cell for 15 hours. Cells were fixed, permeabilized, and stained with EEA1 (Early Endosome Antigen I), pp65, and DAPI. Images were acquired using a 63x oil immersion objective. Scale bars 5 µm. (*N* = 3); representative images are shown.

### UL141 expression can rescue the UL14 growth defect in epithelial cells

Given the structural homology conserved between UL14 and UL141 and the reduced viral infectivity of epithelial cells reported in UL141 deleted viruses (Norris et al., 2025), we wanted to determine if the two related viral proteins have redundant functions in viral tropism. As mentioned, UL141 is truncated in the TB40e backbone used in these studies. So, we wanted to evaluate the impact of restored full length UL141 expression on the growth kinetics of our virus in the absence of UL14. To do this, we generated a panel of mutants using BAC recombineering in the TB40/E background (**Figure 9a**). This panel consists of the previously mentioned UL14-FLAG, ΔUL14, and Repair. In addition, ΔUL14/UL141R was generated by repairing the premature stop codon in the ΔUL14 BAC, restoring UL141 expression and UL14-FLAG/UL141R was generated from the UL14-FLAG BAC where the premature stop codon was also repaired thus allowing expression of both UL14 and UL141. We then performed an extended time course, up to 15 days post infection. Viral output was measured by TCID_50_ for each virus at each timepoint. We observed that the ΔUL14/UL141R and UL14-FLAG/UL141R grew slightly better than the UL14-FLAG virus, lacking UL141 (**Figure 9b**). As expected, the UL14 Repair demonstrated growth kinetics equivalent to the UL14-FLAG virus. Interestingly, the repair of UL141 in the ΔUL14 background was sufficient to rescue the growth defect in comparison to the ΔUL14 virus. This suggests that the functions of UL14 that are undermined in deletion of the viral protein are complemented by UL141 thus demonstrating a level of redundancy in biological functions of UL14 and UL141in terms of efficient epithelial cell infections.

**FIGURE 9:**
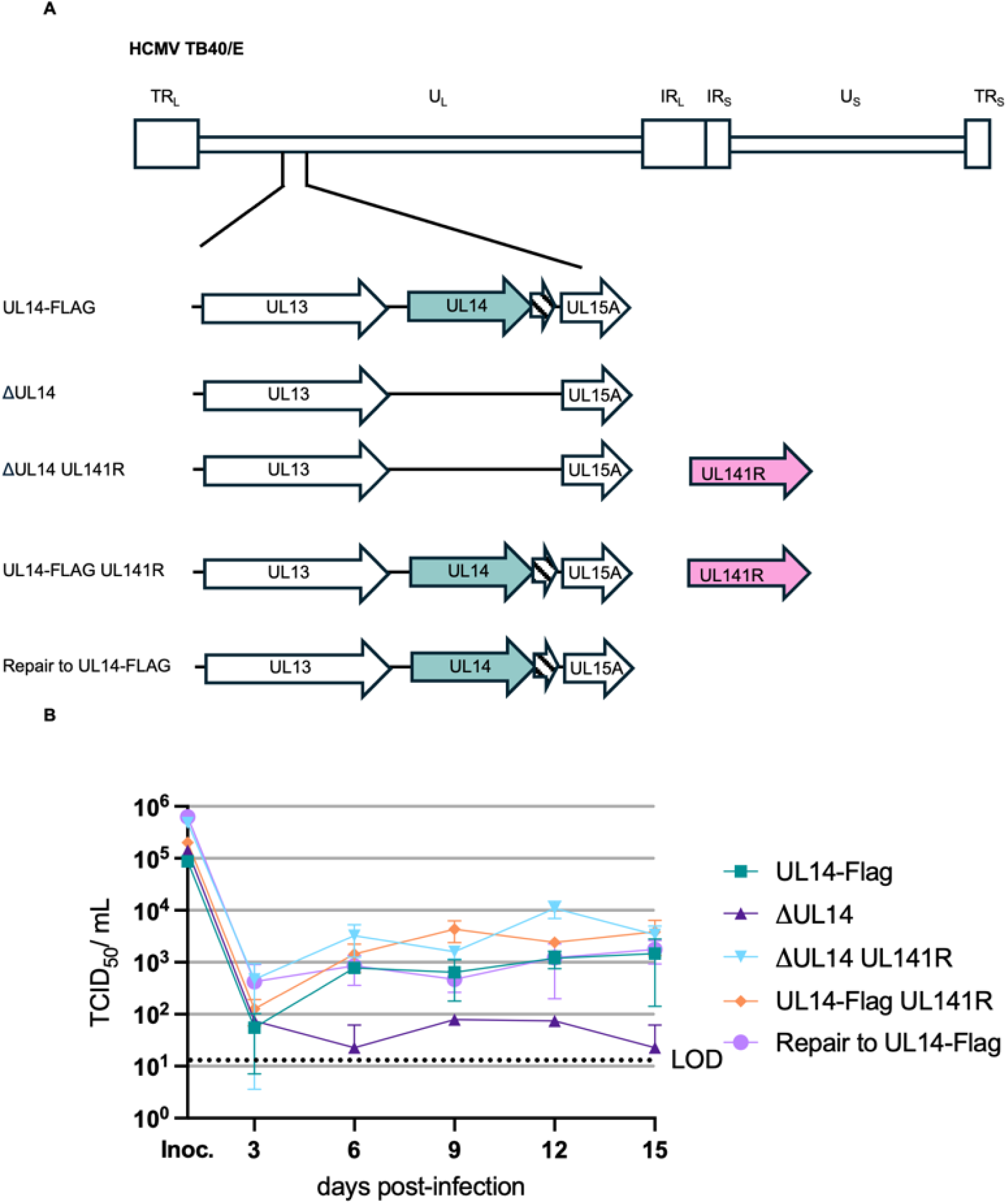
UL141 expression can rescue the UL14 growth defect in epithelial cells. **a**) Schematic of UL14 and UL141 viral mutants generated using BAC recombineering in the TB40/E background. **b)** Epithelial cells (ARPE-19) were infected with the indicated viruses at a multiplicity of infection of 1.0 TCID_50_/cell. Cell associated virus was collected over a time course of 15 days and then viral titers were quantified for each time point by TCID_50_ assays. Mean ± SD is shown (*N=3*).

## DISSCUSSION

Herein, we describe the characterization and biological requirements of UL14 in efficient epithelial cell infections. We report that the UL14 ORF encodes a glycosylated viral protein that is packaged within the virion (**Figure 4a and 4b**), and that this protein is critical for the establishment of infection in epithelial cells (**Figure 5b**). Furthermore, we find that the number of viral genomes delivered per cell is equivalent (**Figure 6c**), however the nuclear delivery of pp65 is diminished and the ΔUL14 virus co-localizes with EEA1 **(Figure 8)**, suggesting that binding and entry of the virus is not impacted, but that the downstream processes including endosome escape are impacted in the absence of UL14. This result may be due to the requirement of an interaction of UL14 with other glycoproteins during the pH-dependent fusion process, including gB and gH, or potentially its interaction with a host receptor. Overall, we have demonstrated a novel role for UL14 as a factor critical in the establishment of infection in epithelial cells, similar to that of its viral homolog UL141.

Given that UL14 is highly conserved amongst laboratory strains that have been highly passaged in fibroblasts, it is reasonable to anticipate that UL14 has additional roles during viral infection that are not addressed in this study. The structural homolog of UL14, HCMV UL141, has been shown to have a role in dampening the innate immune response in addition to its recently identified role in viral entry. Independent of its biological functions within the GATE-3 complex, UL141 has been demonstrated to function in the retention of various receptors and ligands in the endoplasmic reticulum, including the TRAIL death receptors (Nemčovičová et al., 2013; Smith et al., 2013), CD155 (Tomasec et al., 2005), and CD112 (Prod’homme et al., 2010). Although, we were unable to demonstrate a role for UL14 in TRAIL related cell death (data not shown), this may suggest that UL14 and UL141 share partial redundancy in function in terms of viral entry, but that both possess biological functions that are diverse as well.

With the broad cellular tropism demonstrated by HCMV, it is unsurprising that the virus utilizes such a large repertoire of glycoproteins to facilitate its entry. The redundancy demonstrated with UL141 and UL14 leads us to speculate why HCMV would retain both proteins. It is possible that UL14 is structurally more adept for facilitating entry into distinct subsets of epithelial cells, whereas UL141 is geared toward endothelial cell entry, as demonstrated by the Kamil group, where they observed a 5-fold difference in epithelial cells using a virus lacking UL141 compared to endothelial cells where they observed a 1-log difference (Norris et al., 2025). The conservation of both proteins may also be related to their viral interactors, where UL141 binds UL116 and gH to form the GATE-3 complex, and the interacting partners of UL14 remain undefined, but currently under investigation. Follow up studies regarding host interactors of both UL14 and UL141 will need to be conducted to further elucidate their similarities and differences in the context of viral infection.

Overall, our study demonstrates a novel role for a previously uncharacterized viral protein, UL14, which is required for efficient viral infection of epithelial cells. Supporting the notion that the characterization of viral glycoproteins is of critical importance to expand our understanding of viral tropism and spread. Future work aimed at establishing the host receptor and viral interactors will further elucidate the importance of UL14 during natural infection and open the possibilities of the generation of suitable viral targets for the generation of a protective HCMV vaccine.

## MATERIALS AND METHODS

### Cells and viruses

hTert MRC-5 embryonic lung fibroblast (passage 10-30), primary newborn human foreskin fibroblasts (NuFF-1; Global Stem, passage 10-30) and adult retinal pigmented epithelial cells (ARPE-19; ATCC; passage 34-40) were cultured in Dulbecco’s Modified Eagle’s Medium (DMEM), supplemented with 10% fetal bovine serum (FBS), 2mM L-glutamine (523-100p, Cleveland Clinic Media Core), and 100 U/mL penicillin-streptomycin solution (721-100p, Cleveland Clinic Media Core). All cells were maintained at 37°C and 5% CO_2_.

The parental BAC-derived isolate (TB40/e-BAC, clone 4) (Sinzger et al., 2008) was previously engineered to contain SV40 driven mCherry, expressed with IE kinetics, and a UL99 fused GFP, expressed with late kinetics to generate *TB40/E-mCherry-UL99eGFP* (WT) (Nukui et al., 2018). WT was then used to generate the subsequent recombinants using *Galk* BAC recombineering; a UL14 expressing a triple FLAG epitope tag at the C-terminus of the ORF *TB40/E-mCherry-UL99eGFP* UL14-3xFLAG (UL14-FLAG), a deletion of the entire ORF from start to stop codon, using UL14-FLAG as the template *TB40/E-mCherry-UL99eGFP* ΔUL14 (ΔUL14), and then a repair to WT using the ΔUL14 virus as the template *TB40/E-mCherry-UL99eGFP* Repair (Repair). Furthermore, the UL14-FLAG virus and ΔUL14 virus were used as the template to generate a UL141 repair, where the truncated UL141 in TB40/E was restored in each virus, denoted as ΔUL14 UL141R and UL14-FLAG 141R.

As described previously (Warming et al., 2005), the galactokinase gene (*galK*) was amplified using primers designed with 50-bp homology to the flanking regions of interest (Table S1). This product was then purified and electroporated into competent SW105 *Escherichia coli* harboring the*TB40/E-mCherry-UL99eGFP* BAC which were induced by 42°C heat shock for 15 minutes to induce the expression of the *red recombinase* genes. Positive recombinants were then selected for their ability to grow on agar plates in which galactose was the only provided carbon source and confirmed to contain the *galK* insertion using PCR. These positive recombinants were then made electrocompetent and electroporated with the respective duplex oligo or gene block (Table S1) and plated onto 2-deoxy-galactose (2-DOG) minimal plates to select for those BAC constructs that successfully recombineered out the *galK* gene with the targeted template. Positive clones were then picked and sequentially streaked onto minimal media plates containing galactose, then minimal media plates containing 2-deoxy-galactose, and lastly LB+CM plates. Clones that grew on the 2-DOG plates and the LB+CM plates but not the minimal media plates containing galactose were then verified by PCR and Sanger sequencing, or full genome sequencing (Oxford Nanopore Technology).

### Virus propagation and growth assays

Viral BAC DNA isolation, transfection, expansion, and tittering were carried out as previously described (Carter et al., 2025). Briefly, BAC DNA was transfected into hTert MRC5-cells (10cm dish, ∼1 x 10e6 cells), allowed to reach 100% cytopathic effect, and then cell-associated and cell-free virus was isolated. One-tenth of the supernatant was then expanded on naïve NuFF-1 cells (9 15cm dishes, ∼4.5 x 10e7 cells). Cell-associated and cell-free virus was harvested by ultracentrifugation through a 20% sorbitol cushion. The viral pellet was then resuspended in full media containing 1.5% bovine serum albumin, flash frozen with liquid nitrogen, and stored -80°C. Titers for each stock were then calculated using tissue culture infectious dose assay (TCID_50_).

Viral growth kinetics were assessed in NuFF-1 cells by infecting at an MOI of 1 and collecting cell-free virus at 24, 48, 72, 96, and 120 hours post infection. For ARPE-19 cells, viral growth kinetics were assessed by infecting at an MOI of 1 and collecting cell-associated virus at 3, 6, 9, 12, and 15 days post infection. Inoculums at t=0 were also collected and tittered to ensure equal input of virus.

### DNA and protein analysis

For assessment of viral genome delivery cells were infected at an MOI of 1, unbound virus was removed after one hour of incubation and washed with PBS. Four hours post-infection the cells were trypsonized for five minutes, spun down at 500x RCF for 5 minutes and washed three times with PBS. The cell pellet was then lysed in TNE buffer (400 mM NaCl, 10 mM Tris pH 7.5, 10 mM EDTA); the samples were then treated with 40 ug proteinase K and 6.4 ug sodium dodecyl sulfate (SDS) and incubated at 37°C for three hours. The DNA was then extracted using phenol/chloroform followed by a sodium acetate precipitation in the presence of linear acrylamide. Viral and cellular DNA were then quantified by qPCR using primers targeting the UL123 region of the viral genome and MDM2 targeting the human genome, respectively. Genome copy numbers were determined through the employment of a BAC that contains both UL123 and MDM2; this was serially diluted to generate a standard curve of known copy numbers. Samples were run in triplicate using a BioRad CFX 96 Real-Time PCR machine.

For protein analysis cell lysates were harvested in RIPA buffer supplemented with protease and phosphatase inhibitors (Thermo Fisher Scientific Pierce Protease and Phosphatase Inhibitor Mini Tablets, EDTA-free), lysed on ice for 30 minutes and then sonicated via bath sonication for 15 seconds. The protein concentration of each sample was then determined by Bradford assay (Bio-Rad protein assay reagent concentrate), in accordance with manufactures instructions. Samples were subsequently denatured in Lamelli buffer at 95°C for 5 minutes. Protein lysates were then separated by SDS-PAGE and transferred to nitrocellulose membrane using the iBlot semi-dry transfer system, according to manufacturer instructions. Blots were then blocked for 1 hour at room temperature in 5% BSA (Thermo Scientific). Proteins were then detected using the following antibodies: anti-FLAG, clone M2 (Millipore Sigma, 1:7,500); anti-pp65, clone 8A8 (1:500); anti-IE1, clone 1B12 (1:1000); anti-β-tubulin, cat no. 12004165, (BioRad, 1:5,000); goat anti-mouse IgG, HRP conjugated secondary antibodies (CST, 1:5,000).

For protein analysis of cell-free virions, viral particles were isolated using a 20% sorbitol cushion, resuspended in TNE buffer, and protein concentration was determined using the Bradford assay. Samples were then processed for western blotting as described above. Proteins were detected using the following antibodies: anti-pp65, clone 8A8 (1:500); anti-gH, clone AP86 (1:200); anti-gO (1:500); anti-gL (1:500); anti-UL130, clone 5G7(1:500); anti-pp150, clone Cl D (1:500); anti-gB, clone 7.17 (1:500); anti-β-tubulin, cat no. 12004165, (BioRad, 1:5,000); goat anti-mouse IgG, HRP conjugated secondary antibodies (CST, 1:5,000).

### RNA analysis

Cells were lysed in TRIzol (ThermoFisher) at room temperature for five minutes; 1-Bromo-3-chloropane was then added, mixed vigorously, and centrifuged at 12,000xg for 12 minutes for efficient phase separation. The aqueous phase was then used for subsequent RNA precipitation, in the presence of glycogen (Roche). After resuspension in nuclease-free molecular grade water (ThermoFisher), the quantity of RNA was determined using a NanoDrop. RNA was then stored at -80°C or used for downstream processing. cDNA was generated using the TaqMan Reverse Transcription Reagents, according to manufacturer instructions. SYBR green (BioRad) with primers targeting UL14 and GAPDH was used to detect the desired transcripts via RT-qPCR. Samples were run in triplicate in a 96-well format using a BioRad CFX 96 Real-Time PCR machine. UL14 transcripts were normalized to GAPDH.

### IFAs

NuFF-1 cells were seeded in IBIDI treated 8-well chamber slides, incubated overnight and infected the following day at an MOI of 1 and maintained for 72 hours post-infection. ARPE-19 cells were infected at an MOI of 5 and maintained for 15 hours post-infection. Following incubation, cells were fixed with 3.7% paraformaldehyde for 15 minutes, blocked and permeabilized with 0.15% Triton-X and 2% bovine serum albumin (BSA) in PBS. Cells were then stained with the following antibodies diluted in 0.15% Triton-X and 2% BSA in PBS: anti-FLAG (Sigma, 1:500), anti-EEA1 (GeneTex, 1:500), anti- pp65 clone 8A8 (1:10), Alexa Flour 488-conjugated anti-mouse (Jackson ImmunoResearch,1:1,000), Alexa Flour 647- conjugated anti-mouse (Jackson ImmunoResearch,1:1,000), Alexa Flour 647- conjugated anti-rabbit (Jackson ImmunoResearch, 1:1,000), and nuclei were stained using 4’-6’-diamidino-2-phenylindole (DAPI,1:20,000). Cells were then imaged on a Marianas system (3i) consisting of a Ziess Axio Observer 7 equipped with a X-Cite mini+ light source (Excelitas) and a Prime BSI Express CMOS camera (Photometrics) using a 63x oil immersion objective, each image was generated by taking z-stacks of each field with 0.5µm slices with SlideBook software (3i). Images were then exported to ImageJ/Fiji for analysis. Representative images from the midpoint of the cells as determined by nuclear position are shown. Images were scaled and normalized for each channel, merged, and then cropped using ImageJ/Fiji software.

### Lentiviral transduction and expression of UL14

Hek-293T cells were transfected with lipofectamine complexes consisting of 5.6 ug of pLV[Exp]-Puro-EF1A. {UL14-3XFLAG} (Vector Builder), 7.1 ug of p-CMV-VSVG, and 14.2 ug of pDR 8.91 in 60uL of lipofectamine 2000 (Thermo-Fisher). The supernatant was collected at 48 and 72 hours post transfection, filtered with a 0.45-micron filter (Sigma), and concentrated using Lenti-X concentrator (Takara). Concentrated lentivirus was then overlayed onto Nuff-1 cells which were subsequently puromycin selected at 0.55 ug/mL for 2 cell doublings. Whole cell lysates were then collected and analyzed by western blot to confirm protein expression.

### Statistical analysis

Data were analyzed using GraphPad Prism and Excel (Microsoft). The statistical test performed for each experiment is indicated in the figure legends and corresponding text. * P<0.05; ** P<0.01; *** P<0.001; **** P<0.0001.

## Acknowledgements

We thank W.J. Britt (University of Alabama, Birmingham, AL, USA), T. Shenk (Princeton University, Princeton, NJ USA), and G. Chan (SUNY Upstate Medical University, Syracuse, NY, USA) for generously sharing antibodies.

---

**Supplemental Figure 1:**
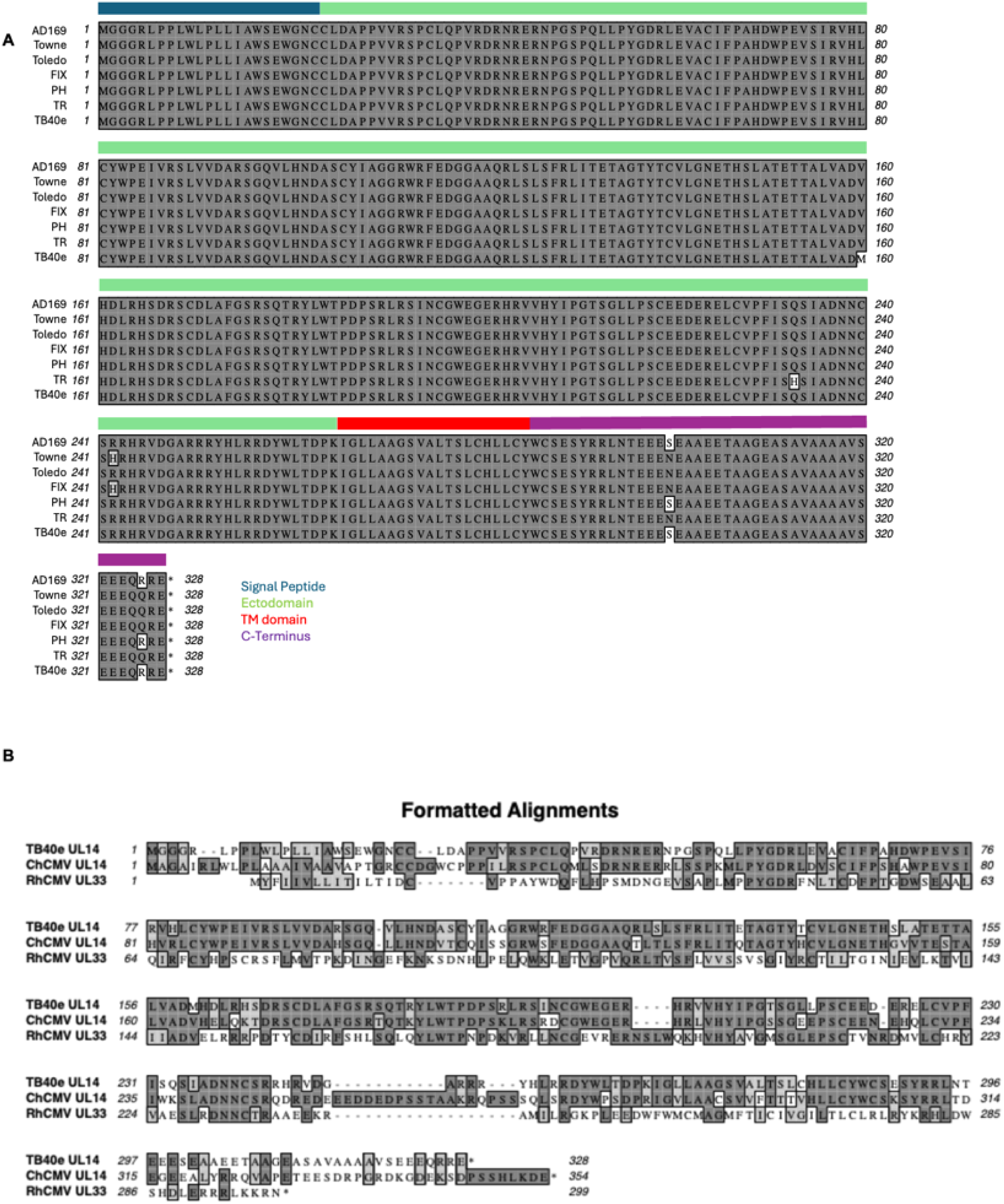
Clustal alignment of UL14. **a**) Amino acid conservation and putative domains of UL14 across AD169, Towne, Toledo, FIX, PH, TR and TB40e are shown. Signal peptide depicted in blue, ectodomain depicted in green, transmembrane domain depicted in red, and the c-terminus depicted in purple. **b**) Amino acid conservation of UL14 from TB40e and primate CMVs; chimpanzee CMV (ChCMV) and rhesus CMV (RhCMV).

**Supplemental Figure 2:**
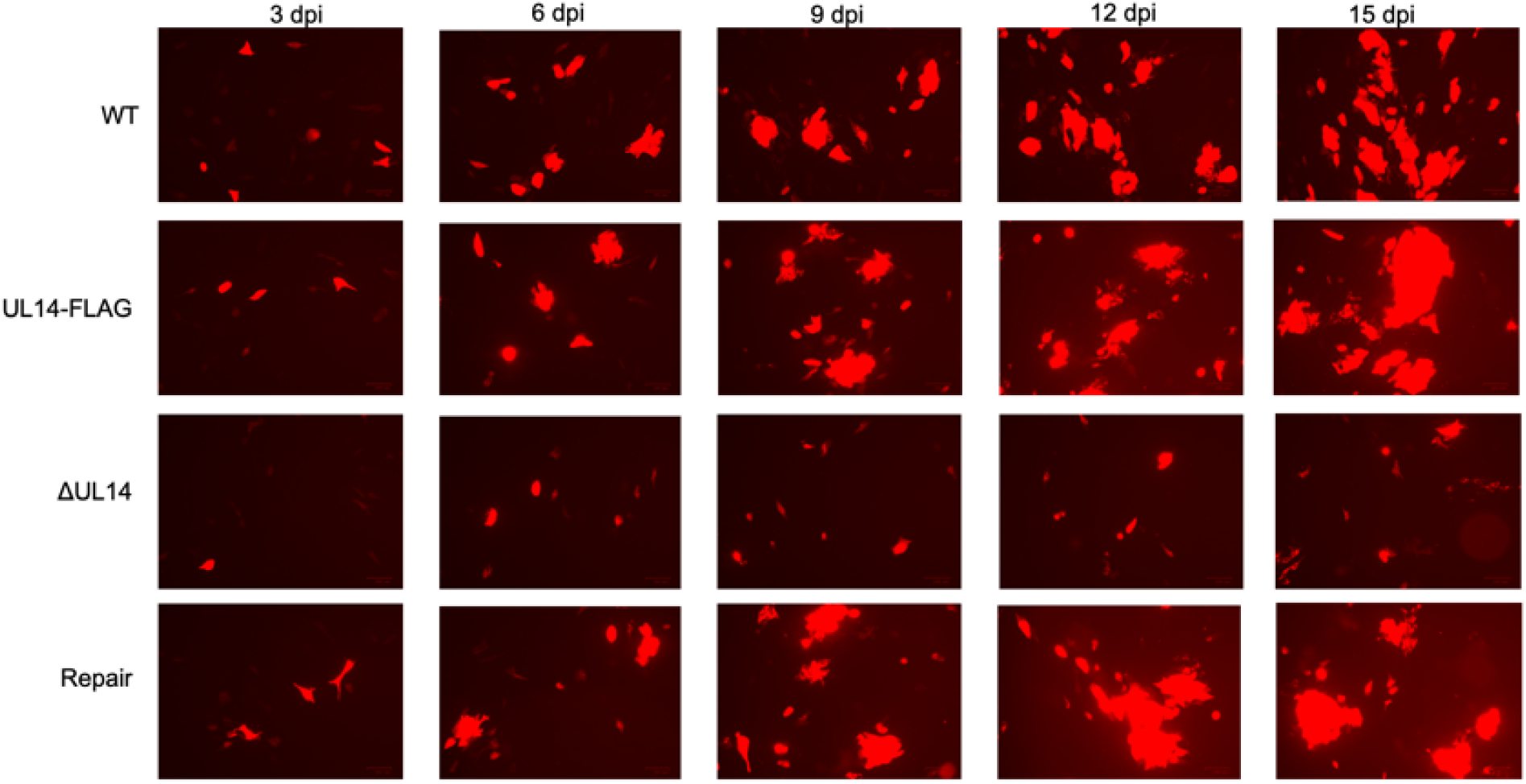
The deletion of UL14 results in a growth defect in epithelial cells. Epithelial cells (ARPE-19) were infected with the indicated viruses at a multiplicity of infection of 1.0 TCID_50_/cell. Images were taken at 3, 6, 9, and 12 days post infection with a BIO-RAD ZOE Fluorescent Cell Imager, where mCherry is being used as a marker of infection. Representative images are shown (*N = 3*).

**Table S1:**
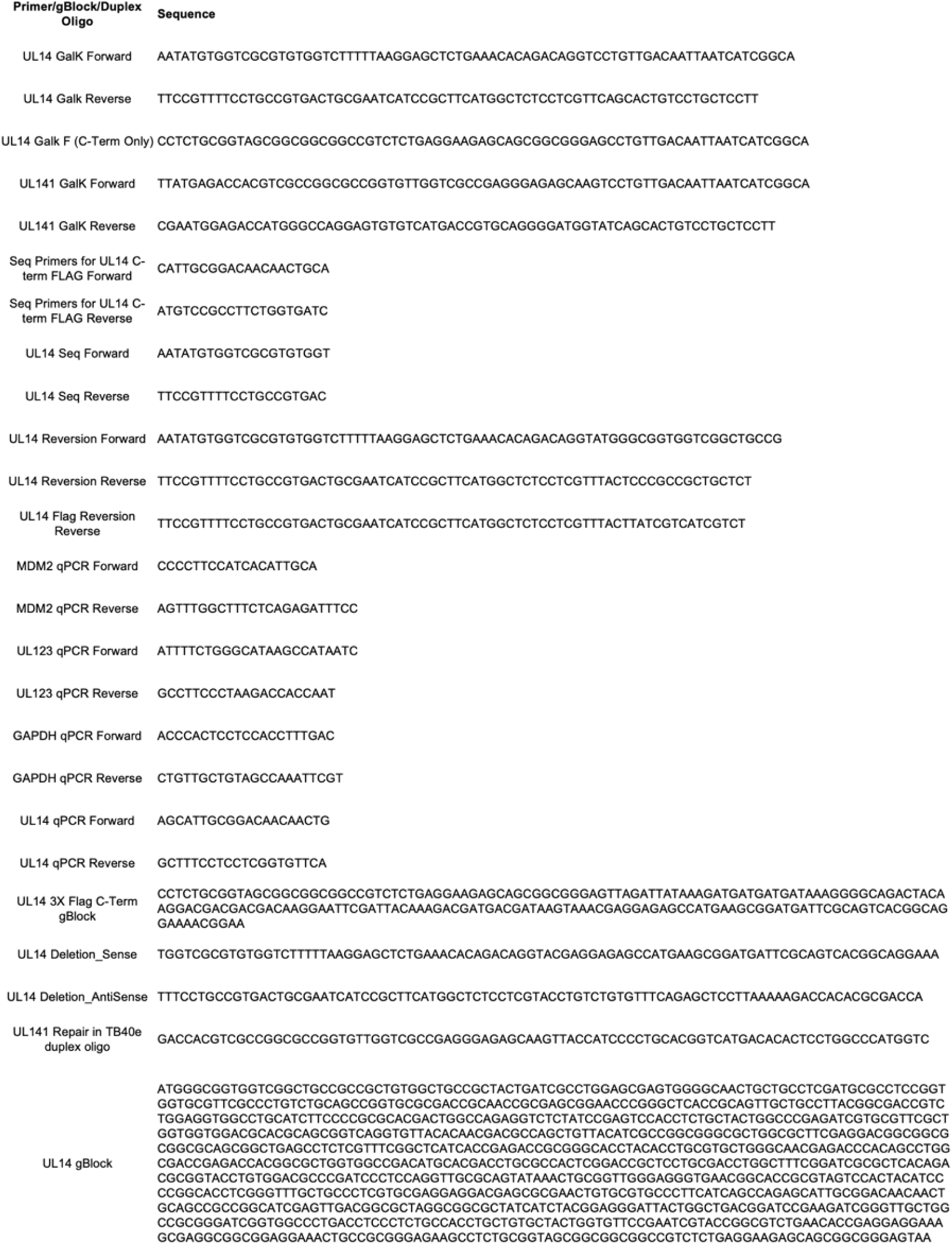

## References

Caló, S., Cortese, M., Ciferri, C., Bruno, L., Gerrein, R., Benucci, B., Monda, G., Gentile, M., Kessler, T., Uematsu, Y., Maione, D., Lilja, A. E., Carfí, A., & Merola, M. (2016). The Human Cytomegalovirus *UL116* Gene Encodes an Envelope Glycoprotein Forming a Complex with gH Independently from gL. Journal of Virology, 90(10), 4926–4938. 10.1128/JVI.02517-15

Carter, M. F., Knight, K., Kayode, Y., & Murphy, E. A. (2025). Generation of a panel of mutants that are resistant to standard of care therapies in a clinically relevant strain of human cytomegalovirus for drug resistance profiling. Antiviral Research, 241, 106237. 10.1016/j.antiviral.2025.106237

Cha, T. A., Tom, E., Kemble, G. W., Duke, G. M., Mocarski, E. S., & Spaete, R. R. (1996). Human cytomegalovirus clinical isolates carry at least 19 genes not found in laboratory strains. Journal of Virology, 70(1), 78–83. 10.1128/jvi.70.1.78-83.1996

Chandramouli, S., Malito, E., Nguyen, T., Luisi, K., Donnarumma, D., Xing, Y., Norais, N., Yu, D., & Carfi, A. (2017). Structural basis for potent antibody-mediated neutralization of human cytomegalovirus. Science Immunology, 2(12). 10.1126/sciimmunol.aan1457

Chesnokova, L. S., & Yurochko, A. D. (2021). Using a Phosphoproteomic Screen to Profile Early Changes During HCMV Infection of Human Monocytes (pp. 233–246). 10.1007/978-1-0716-1111-1_12

Ciferri, C., Chandramouli, S., Donnarumma, D., Nikitin, P. A., Cianfrocco, M. A., Gerrein, R., Feire, A. L., Barnett, S. W., Lilja, A. E., Rappuoli, R., Norais, N., Settembre, E. C., & Carfi, A. (2015). Structural and biochemical studies of HCMV gH/gL/gO and Pentamer reveal mutually exclusive cell entry complexes. Proceedings of the National Academy of Sciences, 112(6), 1767–1772. 10.1073/pnas.1424818112

E, X., Meraner, P., Lu, P., Perreira, J. M., Aker, A. M., McDougall, W. M., Zhuge, R., Chan, G. C., Gerstein, R. M., Caposio, P., Yurochko, A. D., Brass, A. L., & Kowalik, T. F. (2019). OR14I1 is a receptor for the human cytomegalovirus pentameric complex and defines viral epithelial cell tropism. Proceedings of the National Academy of Sciences, 116(14), 7043–7052. 10.1073/pnas.1814850116

Hallgren, J., Tsirigos, K. D., Damgaard Pedersen, M., Juan, J., Armenteros, A., Marcatili, P., Nielsen, H., Krogh, A., & Winther, O. (2022). DeepTMHMM predicts alpha and beta transmembrane proteins using deep neural networks. BioRxiv.

Jumper, J., Evans, R., Pritzel, A., Green, T., Figurnov, M., Ronneberger, O., Tunyasuvunakool, K., Bates, R., Žídek, A., Potapenko, A., Bridgland, A., Meyer, C., Kohl, S. A. A., Ballard, A. J., Cowie, A., Romera-Paredes, B., Nikolov, S., Jain, R., Adler, J., … Hassabis, D. (2021). Highly accurate protein structure prediction with AlphaFold. Nature, 596(7873). 10.1038/s41586-021-03819-2

Kabanova, A., Marcandalli, J., Zhou, T., Bianchi, S., Baxa, U., Tsybovsky, Y., Lilleri, D., Silacci-Fregni, C., Foglierini, M., Fernandez-Rodriguez, B. M., Druz, A., Zhang, B., Geiger, R., Pagani, M., Sallusto, F., Kwong, P. D., Corti, D., Lanzavecchia, A., & Perez, L. (2016). Platelet-derived growth factor-α receptor is the cellular receptor for human cytomegalovirus gHgLgO trimer. Nature Microbiology, 1(8), 16082. 10.1038/nmicrobiol.2016.82

Kari, B., & Gehrz, R. (1993). Structure, composition and heparin binding properties of a human cytomegalovirus glycoprotein complex designated gC-II. Journal of General Virology, 74(2), 255–264. 10.1099/0022-1317-74-2-255

Lantos, P. M., Hoffman, K., Permar, S. R., Jackson, P., Hughes, B. L., Kind, A., & Swamy, G. (2018). Neighborhood Disadvantage is Associated with High Cytomegalovirus Seroprevalence in Pregnancy. Journal of Racial and Ethnic Health Disparities, 5(4), 782–786. 10.1007/s40615-017-0423-4

Martinez-Martin, N., Marcandalli, J., Huang, C. S., Arthur, C. P., Perotti, M., Foglierini, M., Ho, H., Dosey, A. M., Shriver, S., Payandeh, J., Leitner, A., Lanzavecchia, A., Perez, L., & Ciferri, C. (2018). An Unbiased Screen for Human Cytomegalovirus Identifies Neuropilin-2 as a Central Viral Receptor. Cell, 174(5), 1158–1171.e19. 10.1016/j.cell.2018.06.028

Murphy, E., Rigoutsos, I., Shibuya, T., & Shenk, T. E. (2003). Reevaluation of human cytomegalovirus coding potential. Proceedings of the National Academy of Sciences of the United States of America, 100(23). 10.1073/pnas.1735466100

Nemčovičová, I., Benedict, C. A., & Zajonc, D. M. (2013). Structure of Human Cytomegalovirus UL141 Binding to TRAIL-R2 Reveals Novel, Non-canonical Death Receptor Interactions. PLoS Pathogens, 9(3). 10.1371/journal.ppat.1003224

Nguyen, C. C., & Kamil, J. P. (2018). Pathogen at the Gates: Human Cytomegalovirus Entry and Cell Tropism. Viruses, 10(12), 704. 10.3390/v10120704

Norris, M. J., Henderson, L. A., Siddiquey, M. N. A., Yin, J., Yoo, K., Brunel, S., Mor, M., Saphire, E. O., Benedict, C. A., & Kamil, J. P. (2025). The GATE glycoprotein complex enhances human cytomegalovirus entry in endothelial cells. Nature Microbiology, 10(7), 1605–1616. 10.1038/s41564-025-02025-4

Nukui, M., O’connor, C. M., & Murphy, E. A. (2018). The natural flavonoid compound deguelin inhibits HCMV lytic replication within fibroblasts. Viruses, 10(11). 10.3390/v10110614

Permar, S. R., Schleiss, M. R., & Plotkin, S. A. (2025). A vaccine against cytomegalovirus: how close are we? Journal of Clinical Investigation, 135(1). 10.1172/JCI182317

Plotkin, S. A., & Boppana, S. B. (2019). Vaccination against the human cytomegalovirus. Vaccine, 37(50). 10.1016/j.vaccine.2018.02.089

Prod’homme, V., Sugrue, D. M., Stanton, R. J., Nomoto, A., Davies, J., Rickards, C. R., Cochrane, D., Moore, M., Wilkinson, G. W. G., & Tomasec, P. (2010). Human cytomegalovirus UL141 promotes efficient downregulation of the natural killer cell activating ligand CD112. Journal of General Virology, 91(8), 2034–2039. 10.1099/vir.0.021931-0

Rozman, B., Nachshon, A., Levi Samia, R., Lavi, M., Schwartz, M., & Stern-Ginossar, N. (2022). Temporal dynamics of HCMV gene expression in lytic and latent infections. Cell Reports, 39(2), 110653. 10.1016/j.celrep.2022.110653

Scrivano, L., Sinzger, C., Nitschko, H., Koszinowski, U. H., & Adler, B. (2011). HCMV spread and cell tropism are determined by distinct virus populations. PLoS Pathogens, 7(1), e1001256. 10.1371/journal.ppat.1001256

Siddiquey, M. N. A., Schultz, E. P., Yu, Q., Amendola, D., Vezzani, G., Yu, D., Maione, D., Lanchy, J.-M., Ryckman, B. J., Merola, M., & Kamil, J. P. (2021). The Human Cytomegalovirus Protein UL116 Interacts with the Viral Endoplasmic-Reticulum-Resident Glycoprotein UL148 and Promotes the Incorporation of gH/gL Complexes into Virions. Journal of Virology, 95(15). 10.1128/JVI.02207-20

Sinzger, C., Hahn, G., Digel, M., Katona, R., Sampaio, K. L., Messerle, M., Hengel, H., Koszinowski, U., Brune, W., & Adler, B. (2008). Cloning and sequencing of a highly productive, endotheliotropic virus strain derived from human cytomegalovirus TB40/E. Journal of General Virology, 89(2), 359–368. 10.1099/vir.0.83286-0

Smith, W., Tomasec, P., Aicheler, R., Loewendorf, A., Nemčovičová, I., Wang, E. C. Y., Stanton, R. J., MacAuley, M., Norris, P., Willen, L., Ruckova, E., Nomoto, A., Schneider, P., Hahn, G., Zajonc, D. M., Ware, C. F., Wilkinson, G. W. G., & Benedict, C. A. (2013). Human cytomegalovirus glycoprotein UL141 targets the TRAIL death receptors to thwart host innate antiviral defenses. Cell Host and Microbe, 13(3). 10.1016/j.chom.2013.02.003

Song, B., Sheng, X., Justice, J. L., Lum, K. K., Metzger, P. J., Cook, K. C., Kostas, J. C., & Cristea, I. M. (2023a). Intercellular communication within the virus microenvironment affects the susceptibility of cells to secondary viral infections. Science Advances, 9(19). 10.1126/sciadv.adg3433

Song, B., Sheng, X., Justice, J. L., Lum, K. K., Metzger, P. J., Cook, K. C., Kostas, J. C., & Cristea, I. M. (2023b). Intercellular communication within the virus microenvironment affects the susceptibility of cells to secondary viral infections. Science Advances, 9(19). 10.1126/sciadv.adg3433

Teufel, F., Almagro Armenteros, J. J., Johansen, A. R., Gíslason, M. H., Pihl, S. I., Tsirigos, K. D., Winther, O., Brunak, S., von Heijne, G., & Nielsen, H. (2022). SignalP 6.0 predicts all five types of signal peptides using protein language models. Nature Biotechnology, 40(7). 10.1038/s41587-021-01156-3

Tomasec, P., Wang, E. C. Y., Davison, A. J., Vojtesek, B., Armstrong, M., Griffin, C., McSharry, B. P., Morris, R. J., Llewellyn-Lacey, S., Rickards, C., Nomoto, A., Sinzger, C., & Wilkinson, G. W. G. (2005). Downregulation of natural killer cell–activating ligand CD155 by human cytomegalovirus UL141. Nature Immunology, 6(2), 181–188. 10.1038/ni1156

Tsirigos, K. D., Peters, C., Shu, N., Käll, L., & Elofsson, A. (2015). The TOPCONS web server for consensus prediction of membrane protein topology and signal peptides. Nucleic Acids Research, 43(W1). 10.1093/nar/gkv485

van Kempen, M., Kim, S. S., Tumescheit, C., Mirdita, M., Lee, J., Gilchrist, C. L. M., Söding, J., & Steinegger, M. (2024). Fast and accurate protein structure search with Foldseek. Nature Biotechnology, 42(2). 10.1038/s41587-023-01773-0

Vezzani, G., Amendola, D., Yu, D., Chandramouli, S., Frigimelica, E., Maione, D., & Merola, M. (2021). The Human Cytomegalovirus UL116 Glycoprotein Is a Chaperone to Control gH-Based Complexes Levels on Virions. Frontiers in Microbiology, 12. 10.3389/fmicb.2021.630121

Wang, D., & Shenk, T. (2005). Human cytomegalovirus virion protein complex required for epithelial and endothelial cell tropism. Proceedings of the National Academy of Sciences, 102(50), 18153–18158. 10.1073/pnas.0509201102

Warming, S., Costantino, N., Court, D. L., Jenkins, N. A., & Copeland, N. G. (2005). Simple and highly efficient BAC recombineering using galK selection. Nucleic Acids Research, 33(4). 10.1093/nar/gni035

Weekes, M. P., Tomasec, P., Huttlin, E. L., Fielding, C. A., Nusinow, D., Stanton, R. J., Wang, E. C. Y., Aicheler, R., Murrell, I., Wilkinson, G. W. G., Lehner, P. J., & Gygi, S. P. (2014). Quantitative Temporal Viromics: An Approach to Investigate Host-Pathogen Interaction. Cell, 157(6), 1460–1472. 10.1016/j.cell.2014.04.028

Zuhair, M., Smit, G. S. A., Wallis, G., Jabbar, F., Smith, C., Devleesschauwer, B., & Griffiths, P. (2019). Estimation of the worldwide seroprevalence of cytomegalovirus: A systematic review and meta-analysis. In Reviews in Medical Virology (Vol. 29, Number 3). 10.1002/rmv.2034

